# Rapid mechanoenzymatic saccharification of lignocellulosic biomass without bulk water or chemical pre-treatment

**DOI:** 10.1101/2020.03.06.980631

**Authors:** Fabien Hammerer, Shaghayegh Ostadjoo, Karolin Dietrich, Marie-Josée Dumont, Luis F. Del Rio, Tomislav Friščić, Karine Auclair

**Author notes:** Corresponding authors TF and KA.

## Abstract

Lignocellulosic material is an abundant renewable resource with the potential to replace petroleum as a feedstock for the production of fuels and chemicals. The large scale deployment of biomass saccharification is, however, hampered by the necessity to use aggressive reagents and conditions, formation of side-products, and the difficulty to reach elevated monosaccharide concentrations in the crude product. Herein we report the high efficacy of Reactive Aging (or Raging, a technique where enzymatic reaction mixtures, without any bulk aqueous or organic solvent, are treated to multiple cycles of milling and aging) for gram-scale saccharification of raw lignocellulosic biomass samples from different agricultural sources (corn stover, wheat straw, and sugarcane bagasse). The solvent-free enzymatic conversion of lignocellulosic biomass was found to proceed in excellent yields (ca. 90%) at protein loadings as low as 2% w/w, without the need for any prior chemical pre-treatment or high temperatures, to produce highly concentrated (molar) monosaccharides. This crude product of mechanoenzymatic depolymerization is non-toxic to bacteria and can be used as a carbon source for bacterial growth.

## Introduction

With the recognition that use of fossil resources is unsustainable, lignocellulosic biomass has been identified as a main candidate to fulfil the future needs for fuels and basic chemicals.^1–6^ Saccharification of the biopolymer constituents of biomass affords monosaccharides - mainly glucose and xylose - which have been recognized as convenient platforms for the production of valued molecules such as ethanol,^7^ polyhydroxyalkanoates,^8–10^ succinate,^11^ and itaconate.^12^ Despite being produced at an estimated rate of 10^12^ ton per year,^13^ lignocellulosic materials such as agricultural and forestry wastes remain underexploited because of their poor solubility and remarkably recalcitrant nature.^14^ Lignocellulosic materials are composed of cellulose (a linear glucose polymer), hemicellulose (a branched xylose heteropolymer), and lignin (a heterogeneous polyphenolic branched polymer), which are closely intertwined and mostly unaccessible,^11^ posing a persistent challenge for industrial applications.

As a result, chemical hydrolysis of cellulose and hemicellulose to form mono- or oligosaccharides typically requires aggressive chemicals (acids, bases, transition metals)^15–20^ and harsh conditions (temperature, pressure), leading to high energy demands, waste production,^21^ as well as contamination with side reaction products^22–24^ such as furfural, hydroxymethylfurfural (HMF), and acetic acid. Arguably the most important industrial application of glucose, its fermentation by yeasts to produce ethanol, is very sensitive to such impurities.

Whereas biocatalytic processes relying on the action of cellulase and/or hemicellulase enzymes offer a milder, promising alternative for depolymerisation of cellulosic biomass, they are notoriously slow,^25^ and usually require biomass pre-treatment,^26^ again under harsh conditions, to make the biopolymers more accessible to the enzymes.

Another important bottleneck specific to ethanol production from lignocellulosic material is the requirement for a high monosaccharide concentration in the biomass hydrolysate to optimize yeast growth.^7,27–29^ Such concentrations can only be achieved in the presence of high initial amounts of biomass per sample volume (high solid loading), conditions under which the enzymatic saccharification yields are reported to drop significantly. This phenomenon is known as the “solids effect”,^30–32^ and is in part linked to poor homogenization of the mixture and increased inhibition of cellulases by the reaction products.

Our group, and others, have recently demonstrated that enzymes can function surprisingly well in the absence of aqueous or organic solvent, and that their activity can be facilitated by mechanical mixing.^33,34,43,35–42^ This emerging area of research, mechanoenzymology, may provide a solution to the solids effect challenge. In particular, we recently developed a mechanoenzymatic technique (Fig. 1) termed Reactive Aging (RAging), which proceeds in the absence of bulk water, and consists of cycles of alternating periods of brief (minutes) ball milling and longer periods (minutes or hours) of aging,^44,45^ *i.e.* static incubation under controlled conditions. The RAging reactions of cellulases are conducted in the presence of only ca. 10-20 stoichiometric equivalents of water, which acts as a substrate and likely as a reaction lubricant,^69^ leading to a moist solid reaction mixture. This corresponds to solid loadings of 50-100% w/v, which is, to our knowledge, higher than for any previously reported cellulase reactions (typically around 40% w/v).^27,30,32^ At the laboratory scale, our methodology was found to be superior to traditional dilute aqueous reaction mixtures, not only for cellulases,^39,40^ but also for chitinases,^41^ and xylanases.^42^ Although RAging allows direct and efficient depolymerization of microcrystalline cellulose (MCC), as well as cellulose in hay and tree saw dust, without any chemical pre-treatment, further optimization is warranted as earlier reports have been limited to monosaccharide yields not higher than 50%,^40^.

**Figure 1.**
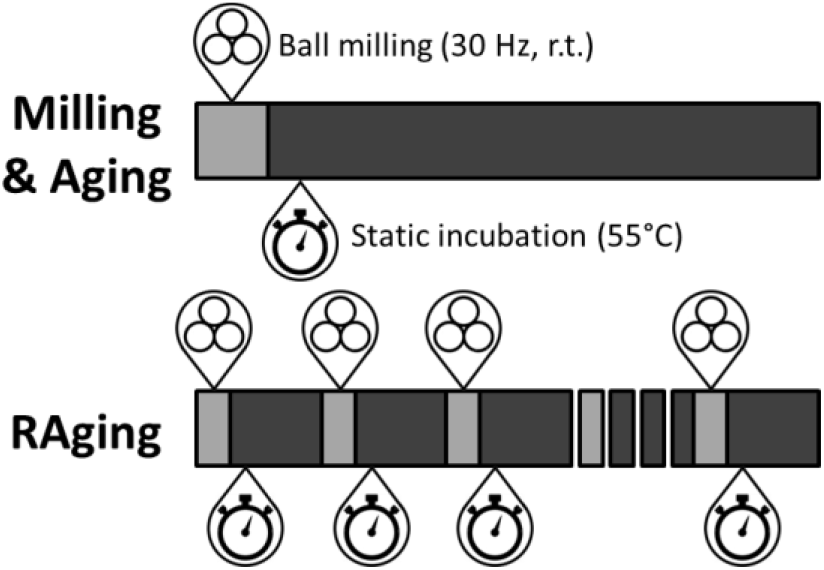
Representation of the mechanoenzymatic processes used in this work: milling followed by aging (top), and RAging (bottom). The commonly accepted symbol for ball milling is used,^46^ while the clock symbol is used to represent aging (static incubation).

We now report the high efficacy, with up to 90% depolymerisation yield of monosaccharides, of cellulases under mechanoenzymatic, water-depleted conditions. This is illustrated in the saccharification of three distinct raw agricultural residues, notably wheat straw (WS), sugarcane bagasse (SB), and corn stover (CS), in a process involving biomass pre-milling and the use of the CTec2 cellulases cocktail (Novozymes). The herein presented methodology generates crude reaction mixtures of monosaccharides in molar concentrations, which we also show can be used directly as a source of carbon in the growth medium of bacterial cultures.

## Results and discussion

### Biomass composition

The experimentally established average compositions (on a dry basis) of the three herein explored agricultural substrates, WS, SB and CS, are presented in Table 1.

**Table 1.** Dry basis composition of the biomass samples used and the maximum extractible monosaccharides.

|  | Biomass composition (%) |  |  | Total extractibles (mmol/g) |
| --- | --- | --- | --- | --- |
|  | Cellulose | Hemicellulose | Others |  |
| Corn Stover (CS) | 33.5 | 23.2 | 43.3 | 3.83 |
| Wheat Straw (WS) | 34.4 | 19.9 | 45.7 | 3.63 |
| Sugarcane Bagasse (SB) | 40.1 | 22.3 | 37.6 | 4.16 |

### Water and protein loading in the mechanoenzymatic reactions

The herein explored RAging mechanoenzymatic reactions proceed with the addition of a small amount of water, which also acts as a substrate. Following the terminology used in context of liquid-assisted mechanochemistry, the amounts of water used correspond to the *η*-parameter, *i.e.* volume of added liquid per sample weight (in μL/mg),^‡^ between 0.5 and 1.5 μL/mg. These conditions meet the previously established regime of mechanochemical liquid-assisted grinding (LAG)^47^ where, as long as *η* is maintained approximately below 2 μL/mg, the reactions can be accelerated or even catalysed^48^ by the presence of a liquid additive, but proceed independent of the relative solubilities of reactants. Due to the absence of solubility limitations typical of reactivity in bulk solvent media,^47^ reactions under LAG conditions are generally considered solvent-free. In our RAging reactions, the added water was completely adsorbed onto the biomass substrate, resulting in reaction mixtures with the appearance of a moist solid. As the reaction progressed, the mixtures were generally found to turn into soft solids with consistencies ranging from those of toothpaste to baking dough.

Another important parameter in enzymatic transformations is enzyme loading, described as the mass of protein per mass of cellulose in the biomass (in mg/g). Protein titers in the commercial *Trichoderma longibrachiatum* cellulases solid stock (13±1% w/w) and in the CTec2 cellulases solution (16±1% w/v) were determined using the Bradford assay. Both of these enzyme preparations contain several cellulases, including exoglucanases, endoglucanases, and β-glucosidases, as well as hemicellulases.

Unless specified otherwise, small-scale milling experiments were performed in 15 mL volume stainless steel jars containing two 7 mm stainless steel balls (1.3 grams each) mounted on a shaker mill operating at 30 Hz. Medium scale milling was accomplished using a 30 mL jar with one 15 mm ball (11.6 grams), both made from stainless steel. The aging part of the reactions was performed by incubation of the milled reaction mixture at 55°C in a closed container.

### Biomass saccharification using *T. longibrachiatum* cellulases

We have previously shown that *T. longibrachiatum* cellulases are superior to the corresponding *Trichoderma reesei* enzymes for cellulose cleavage under mechanochemical conditions.^39^ Thus the latter enzyme preparation was not used here.

After milling raw WS (400 mg; small scale) for 15 min to reduce its size, lyophilized *T. longibrachiatum* cellulases preparation was added (86 mg protein per g cellulose, or 8.6% w/w) together with varying amounts of water. Efficient saccharification of this biomass required a slightly higher liquid/solid ratio (η = 1.34 μL/mL) than MCC (η = 0.9 μL/mL),^39^ leading to 8% digestion of the glycosidic bonds after only 30 min of ball milling, and up to 25% when milling was followed by 3 days of aging at 55°C (Fig. S1; yield estimated using the classical dinitrosalicylic acid, or DNS, method which detects sugar reducing ends^49^). In contrast, RAging of the same reaction mixture, by alternating periods of 5 minutes milling and 55 minutes aging, afforded a yield of 37% in only 12 hours (Fig. 2). Saccharification of SB under the same RAging conditions afforded a 48% yield after 12 hours and 62% yield after 24 hours. When the raw biomass was not pre-milled, the yields went down to 16% and 28% after 12 hours of RAging for WS and SB, respectively.

**Figure 2.**
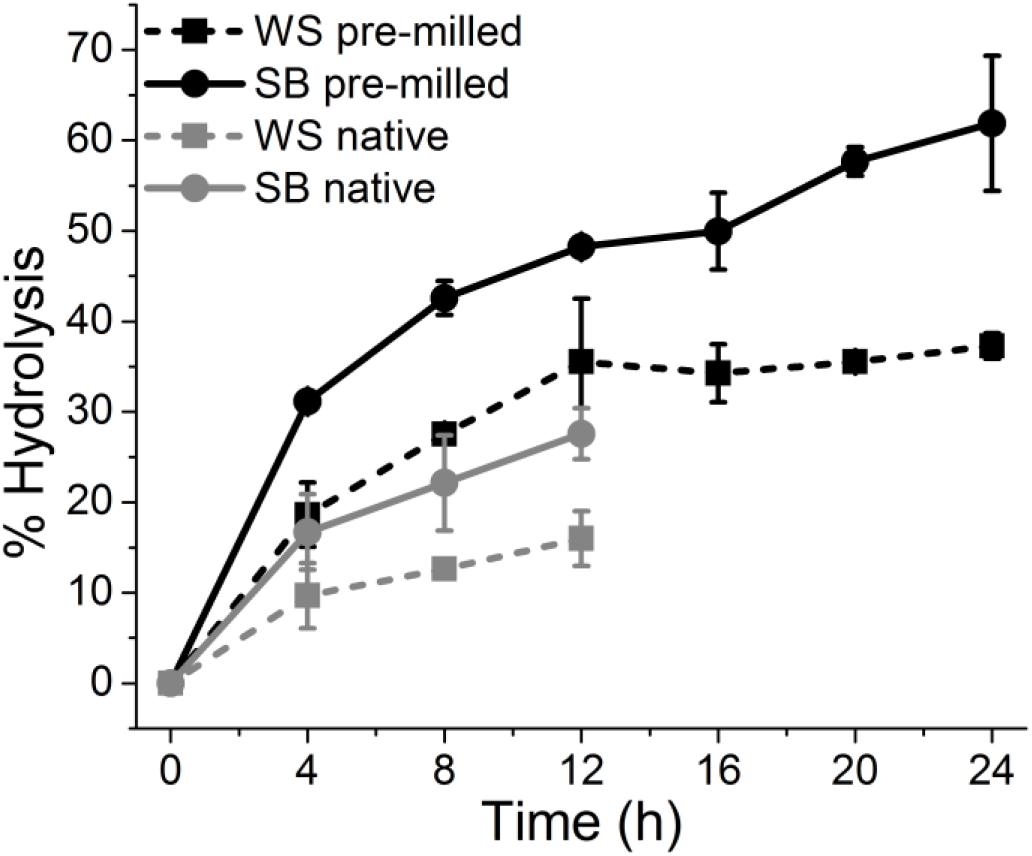
Digestion of native or pre-milled (400 mg, 15 min) WS and SB with the *T. longibrachiatum* enzyme preparation (86 mg/g cellulose) using RAging (cycles of 5 min milling and 55 min aging at 55°C). Yield is DNS-based; error bars are standard deviation from triplicates.

### Biomass pre-milling enhances cellulase activity

Compared to *T. longibrachiatum* cellulases, the commercial CTec2 cellulases blend was found to be more efficient (Fig. S2), and subsequent experiments were performed with this enzyme preparation.

We next assessed the impact of biomass pre-milling on the ensuing activity of CTec2 cellulases. Whereas enzymatic hydrolysis typically relies on a chemical pre-treatment of the biomass,^26^ chemical cellulose saccharification often involves mechanical pre-treatment.^26,50,51^ Thus, CS alone (3 g; medium scale) was treated to various milling durations. The resulting powder was then submitted to a mechanoenzymatic reaction (milling for 30 minutes followed by aging for 3 days) in the presence of CTec2 cellulases (45 mg/g cellulose, 4.5% w/w) at *η* = 1.5 μL/mg. The duration of pre-milling was found to have a large impact on the subsequent mechanoenzymatic transformation (Fig. 3), with a maximum hydrolysis yield of 80±5% obtained when CS was pre-milled for 90 min. Similar results were obtained with the other two biomass substrates, with yields of 73±2% and 75±5% for WS and SB, respectively (Fig. S3). A pre-milling step of 60 minutes duration at a lower CS biomass loading (1.5 g instead of 3 g for the same jar) afforded an even higher yield of 88±2% after enzymatic transformation (Fig. 3).

**Figure 3.**
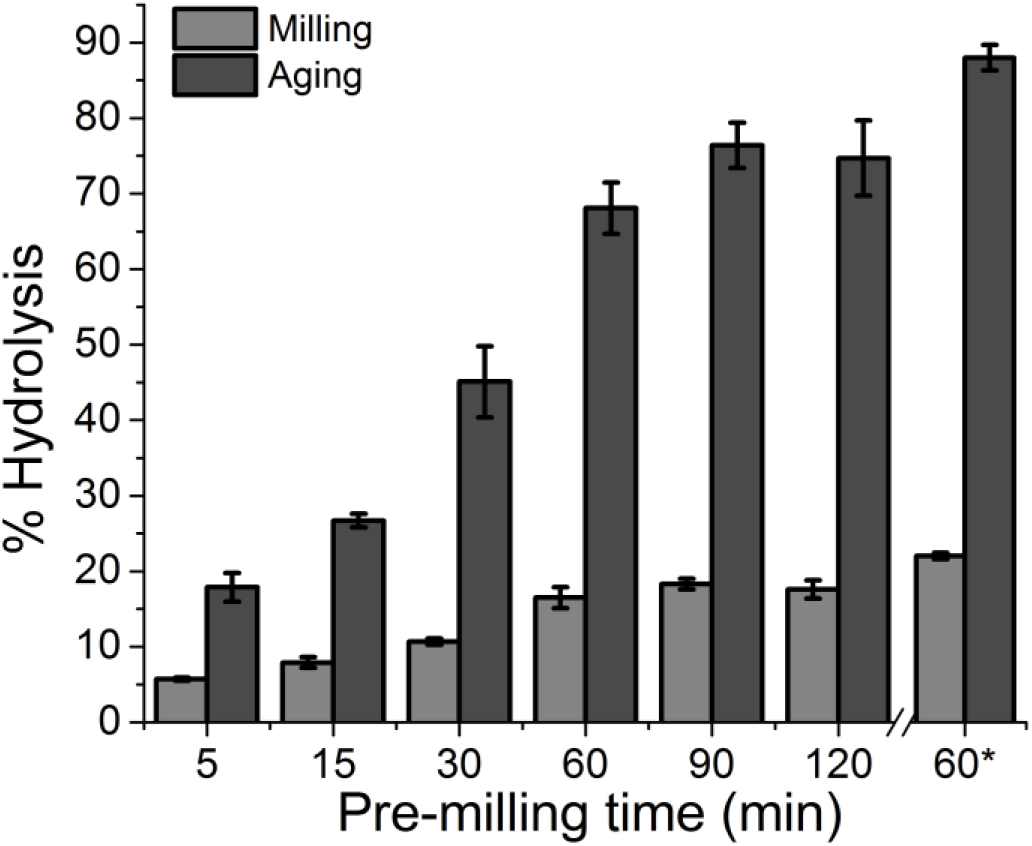
Influence of pre-milling time (3 g in 30 mL jars) on the outcome of milling (30 min) and aging (3 days) reactions of CS with CTec2 enzymes (45 mg/g, *η* = 1.5 μL/mg). Yield is DNS-based; error bars are standard deviation from triplicates. *Jar loading during pre-milling was reduced to 1.5 g.

Besides cellulases, commercial enzymatic preparations may also contain xylanases and other enzymes, indicating that biomass saccharification may lead to a mixture of oligo- and monosaccharides of different sugars, undistinguishable by the DNS assay. The glucose and xylose concentrations of the herein obtained crude reaction mixtures from CS were next measured using a sugar analyser. The glucose concentration was found to be 0.79 M (143 g/L), corresponding to a 57% yield, while the concentration of xylose produced was 0.39 M (60 g/L), corresponding to a yield of 24%. The generally observed variance between the DNS and sugar analysis methods can be explained by the partial digestion of biopolymers into soluble oligosaccharides,^42^ detected by DNS but not with the sugar analyzer.

The crystallinity of CS during the pre-milling step (5-120 min, 3 grams at once) was investigated using powder X-ray diffraction (PXRD) at different time periods, and was found to decrease with milling time (Fig. S5A), with crystalline cellulose (characterized by the Bragg reflection at 2θ of ca. 22°) disappearing within 60 minutes of ball milling. Alone, this reduction in crystallinity cannot account for the increased enzymatic reaction yield, as the maximum conversion is only obtained with 90 minutes of pre-milling. Furthermore, no crystalline cellulose is detected after 60 min of pre-milling, regardless of biomass loading (3 or 1.5 grams; Fig. S5B), while the hydrolysis yield is higher at a 1.5 grams loading (Fig. S4), suggesting that an increase in surface area may also facilitate the enzymatic reaction. In support of this conclusion is the fact that when CS of 30% humidity (significantly less brittle and less efficiently comminuted) is pre-milled for 30 min, the subsequent enzymatic saccharification proceeds only to 25% conversion (Fig. S6), compared to >60% from more brittle, dried CS upon similar treatment. Taken together, these results suggest that pre-milling the biomass enhances cellulases activity as a result of reducing both cellulose crystallinity and increasing substrate surface area.

### Optimization of CTEc2 enzyme activity under milling & aging conditions

Depolymerization of CS by CTec2 cellulases (45 mg/g cellulose) under conditions of milling only, exhibited the usual hyperbolic kinetic profile reported for other mechanoenzymatic reactions^39,41,42^ (Fig. S7), confirming the emerging paradigm that hydrolytic enzymes can easily operate under mechanical agitation. This resilience was further highlighted during the subsequent aging step as shown on Fig. 4. Again, the reaction showed a hyperbolic kinetic profile, with a plateau in conversion appearing only after ca. 20 hours. The initial rate of hydrolysis during aging was measured to be 560±40 mM/h, which is 4 times faster than the initial rate of 130±40 mM/h observed during milling.

**Figure 4:**
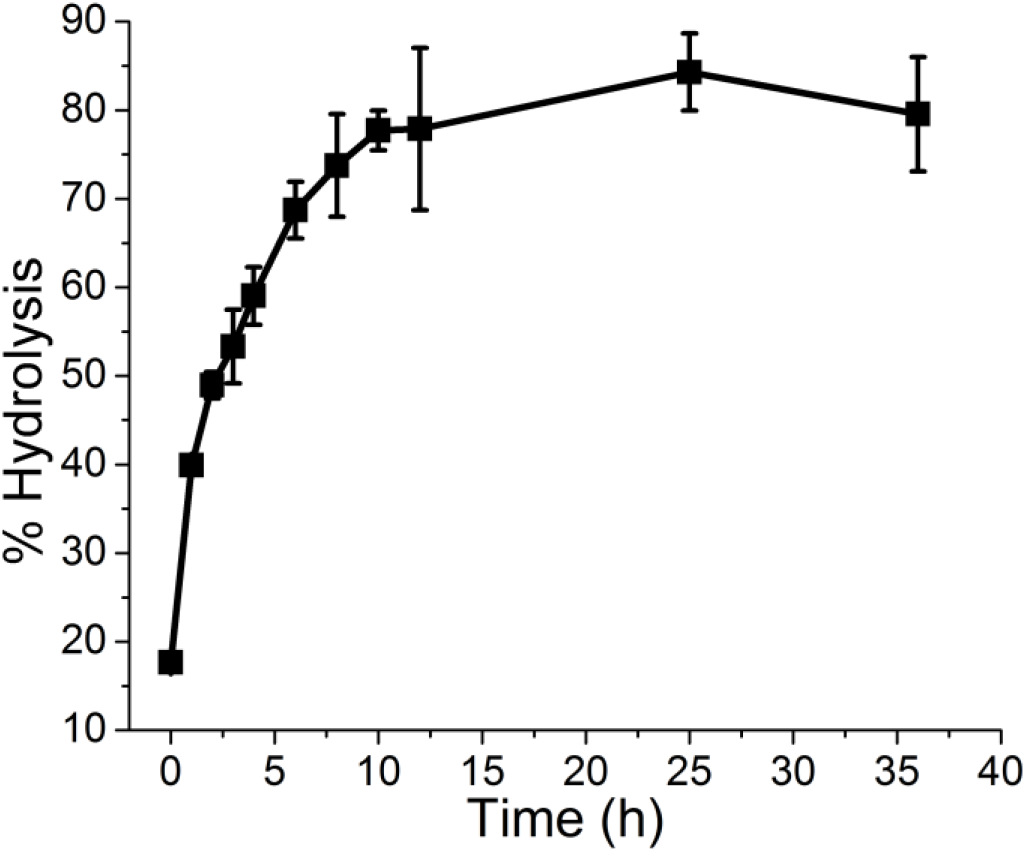
Kinetics of CTec2 enzymes (45 mg/g cellulose, η = 1.35 μL/mg) during aging (after 5 min milling) of a reaction mixture with pre-milled CS (1.5 g, 60 min). The yields were approximated using the DNS assay; error bars are standard deviation for triplicates.

The addition of sodium azide (NaN_3_) – a common antibacterial agent used during enzymatic saccharification reactions – to the enzymatic reaction mixtures (0.04% w/v in the added water) had no significant effect on the reaction yield, even after 3 days of reaction (Fig. S8). In conventional solution-based processes, prolonged enzymatic reactions in dilute buffers are prone to contamination by bacterial or fungal growth. In contrast, the observation that NaN_3_ addition does not affect the outcome of our experiments indicates that solvent-free conditions may not favour microbial growth. Sodium azide was nevertheless used in all subsequent experiments in order to eliminate the possibility of contamination during aging and sample handling.

The structure of biomass exhibits cellulose fibers fully surrounded by hemicellulose, which reduces their accessibility to enzymes. Speculating that this could be a potential limitation of our mechanoenzymatic process, we next explored the use of additional hemicellulase enzymes. The addition of the hemicellulase *Thermomyces lanuginosus* xylanase (1.5 mg/g cellulose) did not improve the yield of cellulose-catalyzed CS hydrolysis (Fig. S9), suggesting that milling and/or the low amount of hemicellulases already present in the CTec2 mixture might be sufficient to make cellulose available for reaction.

CTec2 cellulases exhibited high efficacy during aging, independent of the duration of the milling step (Fig. S10). By milling for only 5 min with the enzyme before aging, we were able to cut the total milling time significantly, without impacting the overall yield.

We also investigated the effect of varying enzyme loading on the efficacy of the mechanoenzymatic depolymerization of CS. Reactions containing 10-50 mg of protein per gram of cellulose were milled for 5 min and aged for 1 or 3 days (Table 2). As is often observed with cellulases in bulk water, higher loadings did not always improve reaction yields. This is usually attributed to limited substrate available at the surface of the biomass.^52,53^ In this case, maximum efficacy was reached at 20 mg/g enzyme loading (or 2% w/w).

**Table 2:** Influence of enzyme loading on milling, and milling & aging reactions. Reaction mixtures contained 400 mg of CS (pre-milled for 60 min at 1.5 g scale) combined with a 600 μL aqueous solution of NaN_3_ (0.04% w/v) and CTec2 cellulases. Mixtures were milled for 5 min at 30 Hz and aged for 1 or 3 days at 55°C.

| Enzyme loading<sup>b</sup> | Reducing sugar yield (%)<sup>a</sup> |  |  |
| --- | --- | --- | --- |
|  | Milling<br>(5 min) | Aging<br>(1 day) | Aging<br>(3 days) |
| 10 | 6.5 ± 0.3 | 61 ± 1 | 71 ± 2 |
| 20 | 12.3 ± 1.2 | 69 ± 5 | 79 ± 2 |
| 30 | 15.1 ± 1.0 | 68 ± 2 | 80 ± 5 |
| 40 | 15.8 ± 0.9 | 71 ± 4 | 79 ± 6 |
| 50 | 14.8 ± 1.1 | 64 ± 2 | 79 ± 3 |

a) DNS-based. Error is standard deviation from triplicates. b) Reported in mg protein/g cellulose, corresponding to 1-5% w/w.

Remarkably, the reaction can be conveniently scaled up from 400 mg to 1.5 g, affording 81±5%, 73±3% and 83.1±0.8% after 3 days of aging for CS, WS and SB respectively, similar to smaller scale reactions (Fig. S11).

### Enhanced reactivity under RAging conditions

We further looked to accelerate biomass hydrolysis by CTec2 using RAging. Pre-milled CS was hydrolysed enzymatically via multiple 1-hour cycles, each consisting of 5 min of milling and 55 min of aging. Saccharification reached 83% after only twelve cycles (Fig. 5, Table 3). Even though the yield (as measured by the DNS assay) was roughly the same as with milling only once followed by 3 days of aging, it was reached much faster (6 times). Furthermore, detailed analysis revealed a glucose content ~30% higher with RAging than through a combination of a single milling step followed by aging, and a slightly improved xylose content. Reproducibility was also significantly improved (comparing Tables 2 and 3).

Enzymatic hydrolysis of WS and SB under the same RAging conditions proceeded in 70% and 76% yields, respectively. These values, however, rose to 84 or 86%% if the 12-hour RAging period was followed by an additional 12 hours of aging (Table 3).

**Table 3:** RAging pre-milled biomass. Reaction mixtures contained 400 mg pre-milled substrate (1.5 g, 60 min) combined with 600 μL of an aqueous solution of NaN_3_ (0.04% w/v) and CTec2 cellulases (45 mg protein/g cellulose). Reactions were submitted to 12 cycles of 5 min milling at 30 Hz and 55 min aging at 55°C followed by another 12 h of aging at 55°C.

| Substrate | RAging (12 h) |  |  |  |  |  | RAging (12 h) + Aging (12 h) |  |
| --- | --- | --- | --- | --- | --- | --- | --- | --- |
|  | Reducing sugars<sup>a</sup> |  | Glucose<sup>b</sup> |  | Xylose<sup>b</sup> |  | Reducing sugars |  |
|  | Yield | Final concentration (M) | Yield | Final concentration (M) | Yield | Final concentration (M) | Yield | Final concentration (M) |
| CS | 83 ± 3% | 2.09 ± 0.06 | 77±1 % | 1.05±0.01 | 39±2 % | 0.4±0.1 | 86 ± 2% | 2.16 ± 0.03 |
| WS | 70 ± 1% | 1.61 ± 0.01 | 65±4 % | 0.85±0.05 | 39±3 % | 0.4±0.1 | 84 ± 4% | 1.90 ± 0.09 |
| SB | 76 ± 4% | 1.96 ± 0.09 | 66±2 % | 1.0±0.2 | 37±5 % | 0.3±0.1 | 84 ± 4% | 2.14 ± 0.01 |

a) Based on the DNS assay. b) Measured by sugar analysis. Error is the standard deviation for triplicates.

**Figure 5:**
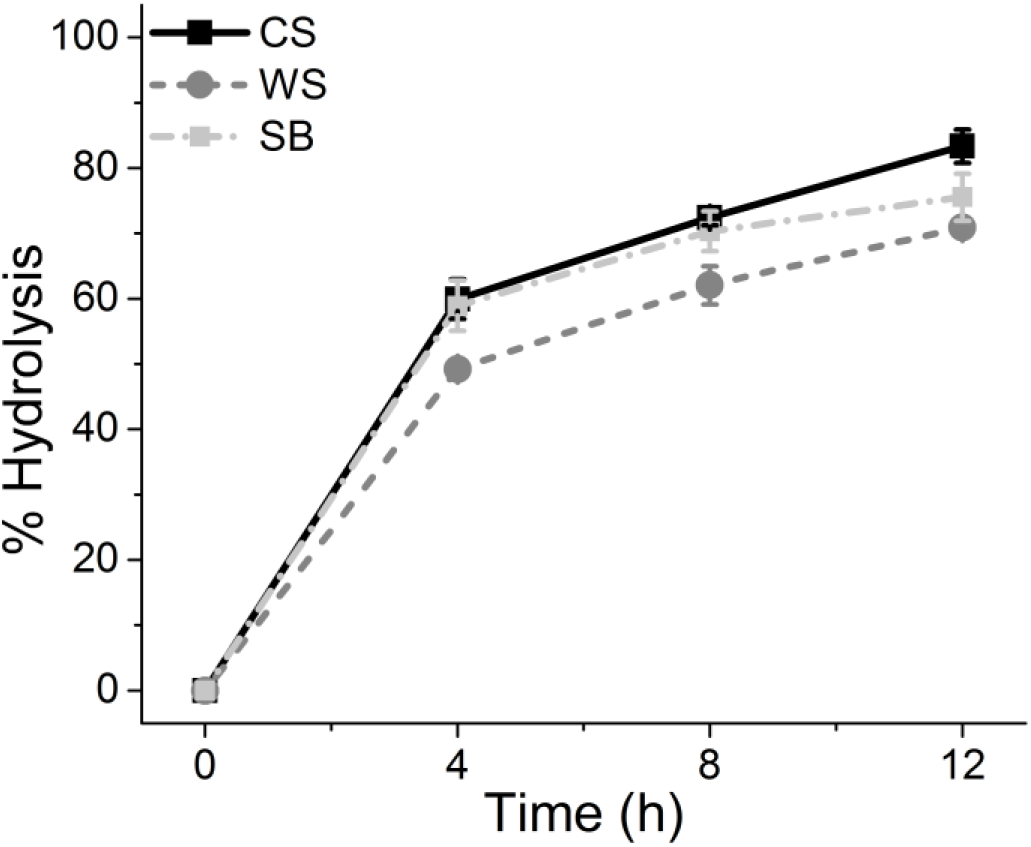
RAging pre-milled CS (full black line), WS (dashed, dark grey), and SB (dashed, light grey). Reaction mixtures contained biomass (400 mg, pre-milled for 60 min) combined with 600 μL of an aqueous solution containing NaN_3_ (0.04% w/v) and CTec2 (45 mg protein/g cellulose). Mixtures were submitted to 12 cycles of 5 min milling and 55 min aging. Yields are approximated with the DNS assay. Error bars are standard deviations from triplicates.

Compared to conventional slurry or solution processes which require a harsh pre-treatment, RAging does not require any chemical pre-treatment, while leading to higher reaction rates (Tables S1, S2).^29,32,62,54–61^ In addition to providing the crude product with the highest reported monosaccharide concentration (second highest for glucose alone), the space-time yield (mass of sugar produced per litre of reaction per hour, see Tables S1 and S2) of our enzymatic RAging process is at least twice higher than that of any other reported method (Tables S1 and S2).

### Crude RAging products as a carbon source for bacterial growth

Next, we evaluated the potential of our biomass hydrolysates for use as the carbon source in bacterial growth media. As highlighted in our previous report,^39^ sugars can be conveniently recovered from crude reaction mixtures by centrifugation. Thus, from crude RAging reaction mixtures, the resulting sugar-rich supernatant was diluted to ca. 0.3% w/v based on monosaccharides^§^, supplemented with standard mineral salts (carbon-free), and directly used as a bacterial culture medium using standard protocols. Both *Escherichia coli* and *Salmonella enterica* ser. Typhimurium were found to proliferate equally well on agar gels derived from either standard lysogeny broth (LB) or media prepared from crude saccharification reaction mixtures from either CS, WS, or SB (Fig S12).

We next looked at planktonic growth of a strain of *Paraburkholderia sacchari*^10^ known to metabolize both glucose and xylose, and to produce polyhydroxybutyrate, a highly valued polymer and important candidate for large scale deployment of biodegradable plastics.^9^ The hydrolysate resulting from milling & aging SB was used as the sole carbon source of the growth medium, adjusted to final concentrations of 11.6 g/L glucose and 5.7 g/L xylose. Bacteria were inoculated in this medium, and after 24 hours of incubation, they proliferated to 6.9±0.3 g/L cell dry mass (∆CDM), which is higher than for a control experiment containing 20 g/L of pure glucose instead of biomass hydrolysate (Fig. 6A). While the glucose consumption was similar in both experiments (around 11 g/L) the hydrolysate allowed the consumption of an extra 3.5 g/L of xylose (Fig. 6B). These experiments demonstrate that the crude products of our mechanoenzymatic reactions can be used directly as the only carbon source in fermentation processes without toxicity.

**Figure 6:**
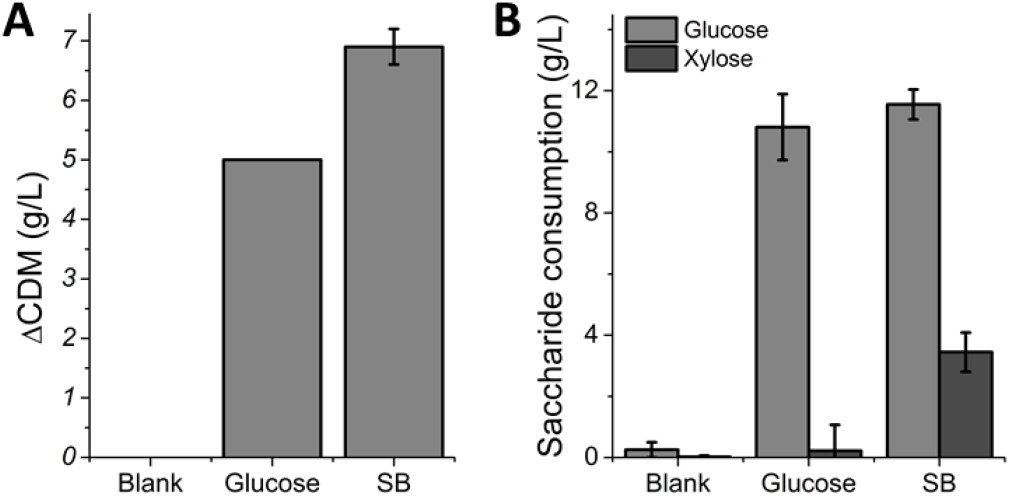
A) *P. sacchari* growth in a culture medium either without a carbon source (blank), prepared from commercial glucose (20 g/L), or from a RAging-derived SB hydrolysate (final titer: 11.6 g/L glucose and 5.7 g/L xylose). B) Saccharide consumption after 24 hours of *P. sacchari* growth using commercial glucose or SB hydrolysate as the main carbon source.

## Conclusions

We report here that both RAging, as well as aging after a brief period of mechanochemical activation, enable the enzymatic breakdown of cellulose and xylan in the absence of bulk water, directly from different types of biomass, without any need for chemical pre-treatment. Simply pre-milling of the raw lignocellulosic material for 60-90 min in order to obtain a fine powder, before enzyme addition, was sufficient to ensure subsequent mechanoenzymatic saccharification yields of ca. 90% within 12 to 24 hours on a gram scale. Moreover, the enzymatic depolymerization of all three biomass substrates proceeded to yield molar-level concentrations of glucose and xylose monosaccharides. After separation from the solids, the crude sugars were efficiently used as a carbon source for bacterial growth on agar gels or in a bioreactor, demonstrating their low toxicity and biocompatibility.

Biocatalysis is appreciated for its selectivity, mild conditions, low toxicity, and catalyst renewability. The growing field of mechanoenzymology provides new exciting opportunities for biocatalytic transformations. Not only were enzymes reported to tolerate mechanical stress,^33–42^ but they were also found to remain active in moist solid mixtures.^43^ Static incubation of enzymatic reaction mixtures without bulk water may provide a better mimic of the natural environment of enzymes, and especially so for enzymes that are secreted by soil microorganisms that thrive on moist surfaces rather than dilute aqueous solutions.^67,68^ By minimizing the total volume of the reaction mixture, the herein presented mechanochemically-activated enzymatic processes greatly facilitate handling and mixing, and curtail waste associated with processing and depolymerization of polysaccharide biomass. Furthermore, as highlighted in this study, mechanoenzymology avoids solubility issues and the solids effect which normally impairs enzymatic saccharification of lignocellulosic materials at high solid loading. Consequently, we believe that this non-traditional way of using enzymes should find broad application as a cleaner, simpler and more efficient route for converting biomass into well-defined small molecules, without requiring bulk solvents, strong mechanical impact or high temperatures.

## Methods (see footnotes for references)

### Material and general methods

Wheat straw (WS) and sugarcane bagasse (SB) were kindly provided by Iogen Corporation (Ottawa, Canada), and corn stover (CS) was from POET (Sioux Falls, SD, USA). Cellulases from *Trichoderma longibrachiatum* (G4423), xylanase from *Thermomyces lanuginosus*, and the CTec2 preparation (SAE0020) were purchased from Sigma-Aldrich. 3,5-Dinitrosalicylic acid, potassium sodium tartrate tetrahydrate, and sodium azide were purchased from Millipore Sigma (Oakville, ON, Canada). Microcrystalline cellulose (MCC) was obtained from Sigma Aldrich (Saint-Louis). Water was from a MilliQ system with a specific resistance of 18.2 MΩcm at 25°C.

Mechanochemical reactions were performed in SmartSnap stainless steel jars (15 or 30 mL) from FormTech Scientific, containing stainless steel balls (7 or 15 mm in diameter) set up on a FTS1000 shaker mill from FormTech Scientific. Aging was performed in an IsoTemp vented oven from Fischer Scientific set at 55°C.

All experiments were performed at least in triplicates, and presented as the average with the standard deviation.

### Biomass preparation and handling

As measured using the standard NREL protocol,^1^ WS and SB had water contents of 6.1% and 6.4% w/w respectively and were used as is. CS was noticeably more humid and partially mouldy, with a water content of roughly 30%. To increase reproducibility, the CS sample was dried at 75°C in a vented oven for several days to a water content of 1.8% w/w, and stored at r.t. None of the substrates were washed before use.

### Reaction analysis and yield determination

Reaction yields were first estimated using the classical dinitrosalicylic acid (DNS) method.^2^ DNS reacts with the reducing end of sugars and does not discriminate between them. Since both enzyme cocktails possess xylanase activity, unless otherwise noted, the yields presented in this study are for hydrolysis of holocellulose i.e. cellulose and hemicellulose combined. Detailed sugar analysis was performed on key samples to determine specific glucose concentrations and yields. The DNS reagent solution was prepared by mixing 3,5-dinitrosalicylic acid (1 g) in deionized water (50 mL), before addition of sodium potassium tartrate tetrahydrate in small portions (total 30 g). An aqueous solution of sodium hydroxide (20 mL of 2 M) was next added. The mixture turned from transparent to an intense yellow color. The volume was adjusted to 100 mL with water and the solution was filtered through cotton. This reagent solution was stored at 4°C in an inactinic container for up to one month.

The frozen reaction mixture aliquots to be analyzed were thawed on ice. Ice cold water was added to obtain a 10 mg/mL suspension of initial biomass weight. The samples were incubated for 30 min at 100°C to inactivate the enzymes. The samples were returned to room temperature and aggregates were broken down using a spatula. The suspensions were then centrifuged for 5 min at 21,100 × *g* and the supernantant was kept for analysis.

The standard procedure for the DNS assay^2^ was followed with a few modifications. In a 1.5 mL microtube, the supernatant of the reaction mixture (50 μL) was diluted in deionized water (150 μL) and the DNS reagent solution (100 μL) was added, before vortexing for 2 seconds, and incubating for 5 minutes at 100°C. After cooling down to room temperature, a portion of the sample (200 μL) was introduced in the well of a 96-well microtiter plate. Absorption at 540 nm was measured using a SpectraMax i3x from Molecular Device with PathCheck enabled. For each new DNS batches, a calibration curve was plotted from analysis of freshly prepared glucose solutions of known concentrations. The reagent was found to be stable for several weeks. The amount of carbohydrates present in the commercial enzymatic mixture was measured and subtracted from the results.

Due to the polydispersity of cellulose, the heteropolymeric nature of hemicellulose, and the presence of both cellulases and hemicellulases in the enzyme mixtures used, yields were calculated on a dry mass basis by approximating cellulose and hemicellulose to linear infinite chains of linked glucose or xylose, respectively.

### Small scale (15 mL jars) pre-milling of biomass

The biomass substrate (400 mg) was introduced in a 15 mL stainless steel jar containing two 7 mm stainless steel balls and milled at 30 Hz for the desired duration. The resulting powder was either used directly or stored in a closed glass vial under ambient conditions.

### Large scale (30 mL jars) pre-milling of biomass

The biomass substrate (1.5 g) was introduced in a 30 mL stainless steel jar containing one 15 mm stainless steel ball and milled for 1 min at 30 Hz. Additional substrate was added if needed to reach the desired final mass (up to 3 g) and with further milling for the desired time (5 to 90 min). The resulting powder was recovered and stored at room temperature in a closed glass vial.

### Enzymatic reactions under milling and aging conditions

Unless otherwise noted, all milling reactions were performed at a 400 mg (small) scale in a 15 mL stainless steel jar containing two 7 mm stainless steel balls. The raw or pre-milled biomass was combined with the *T. longibrachiatum* solid enzyme preparation and the desired amount of water, or with a solution of CTec2 diluted in the appropriate volume of water so that no additional water needs to be added, with or without sodium azide (0.04% w/v). The reaction mixtures were milled at 30 Hz for the desired duration (5-60 min). Aliquots (10-20 mg) were collected, weighed precisely, and frozen until analysis. Aging was accomplished at 55°C for the desired duration (1 h to 3 days) in a closed container inside a vented oven. Aliquots (10-20 mg) were collected, weighed precisely, and frozen until analysis.

### Enzymatic reactions under RAging conditions

All RAging reactions were performed at a 400 mg (small) scale in a 14 mL Teflon jar containing two 7 mm stainless steel balls. The biomass substrate (pre-milled or not) was combined with the *T. longibrachiatum* solid enzyme preparation and the desired amount of water, or with a solution of CTec2 diluted in the appropriate volume of water so that no additional water needs to be added, with or without sodium azide (0.04% w/v). The mixture was milled at 30 Hz for 5 min and transferred to a vented oven set at 55°C for 55 min with the jar kept closed. After this incubation period, the jar was taken out, the mixture milled again for 5 min and aged for 55 min. The jar was submitted to a total of 12 cycles. Aliquots (10-20 mg) were collected after 4, 8 and 12 h, weighed precisely, and frozen until analysis. In some cases, aging was continued uninterrupted for another 1 h to 3 days. Aliquots (10-20 mg) were collected, weighed precisely, and frozen until analysis.

### Reactions with wet CS biomass for comparison

Corn stover (400 mg dried or 570 mg wet to account for 30% humidity) was introduced in a 15 mL stainless steel jar containing two 7 mm stainless steel balls and milled for 30 min at 30 Hz. The *T. longibrachiatum* enzyme preparation (final enzyme loading of 45 mg/g cellulose) was added followed by water (600 or 430 μL for dried or wet biomass respectively; no sodium azide was used for this experiment). The mixture was milled for 30 min at 30 Hz and aged for 3 days at 55°C. The samples were collected, treated, and analyzed as described above.

### Xylanase addition

Corn stover (1.5 g) was pre-milled for 60 min following the protocol described above. The resulting powder (400 mg) was introduced in a 15 mL stainless steel jar containing two 7 mm stainless steel balls and combined with a xylanase preparation from *Thermomyces lanuginosus* (50 mg, 1.5 mg/g cellulose) and a freshly prepared CTec2 solution in water (final enzyme loading 45 mg/g cellulose). The mixture was milled for 30 min at 30 Hz and aged for 3 days at 55°C. The samples were collected, treated, and analyzed as described above.

### Larger scale enzymatic reactions

The pre-milled biomass substrate (1.5 g) was introduced in a 30 mL stainless steel jars containing one 15 mm stainless steel ball. A freshly prepared CTec2 solution (2.25 mL) in aqueous sodium azide 0.04% w/v was added. The mixture was milled for 5 min at 30 Hz. An aliquot (10-20 mg) was collected, weighed precisely, and frozen at −20°C. The rest of the mixture was transferred to a glass vial and aged for 3 days at 55°C. An aliquot (10-20 mg) was collected, weighed precisely, and frozen until analysis. The remaining hydrolysate was stored at −20°C to be later used for bacterial growth and fermentation experiments.

### Analysis of corn stover (CS) crystallinity after pre-milling

X-Ray diffraction (PXRD) patterns of pre-milled CS (1.5 or 3 g in a 30 mL stainless steel jar for 5-90 min) were collected at room temperature on a Bruker D8 Discovery instrument equipped with a LYNXEYE XE-T detector (1D mode), using nickel-filtered Cu*K*_α_ (*λ* = 0.154056 Å) radiation.

### Sugar analysis

The monosaccharides were analyzed in a sugar analyzer (YSI 2900, Yellow Spring Instruments). Crude reaction mixtures were vortexed for 10 seconds, and an aliquot (400 μL) was centrifuged for 5 min at 9400 × *g* before analysis. For the RAging reaction mixtures, a sample was collected (ca. 100 mg), extracted in distilled water (10 mL) by vortexing for 30 seconds. Next, an aliquot (1 mL) was centrifuged for 5 min at 9400 × *g*, and the supernatant was separated and analyzed. This analyzer uses glucose and xylose specific membranes, and calibrates itself automatically with standard glucose (2.5 g/L) and xylose (20 g/L) solutions. The pH is kept at 7.0 by using a mono and dibasic sodium phosphate buffer (YSI 2357, 0.1 M, pH 7.0)

### Agar plates from biomass hydrolysates for the culture of *E. coli* and *S. Typhimurium*

Growth media were prepared according to a protocol by Causey *et al.*^3^ and contained: 3.5 g of KH_2_PO_4_, 5.0 g of K_2_HPO_4_, 3.5 g of (NH_4_)_2_HPO_4_, 0.25 g of MgSO_4_.7H_2_O, 15 mg of CaCl_2_.2H_2_O, 0.5 mg of thiamine chloride, and 1 mL of trace elements solution per liter. The trace elements solution was prepared in 0.1 M HCl and contained: 1.6 g of FeCl_3_, 0.2 g of CoCl_2_.6H_2_O, 0.1 g of CuCl_2_, 0.2 g of ZnCl_2_.4H_2_O, 0.2 g of NaMoO_4_, and 0.05 g of H_3_BO_3_ per liter.

Hydrolysates from aging reactions (21 g containing roughly 4.3 g monosaccharides) from all three biomass substrates (CS, WS, SB) were thawed, suspended in growth medium (20 mL), and transferred to 40 mL centrifuge tubes. The suspensions were centrifuged at 20,000 × *g* for 5 min. The supernatant was collected. The pellet was further extracted by re-suspension in growth medium (20 mL), and breaking down the aggregates with a spatula. After centrifugation (5 min at 20,000 × *g*), both supernatants were combined. The total volume was adjusted to 140 mL with growth medium to afford ca. 3% w/v monosaccharides. These concentrated sugar solutions were partitioned into 10 mL tubes and stored at −20°C until needed.

When needed, a sugar solution (10 mL) was thawed and combined with 90 mL of growth medium for a final concentration of 0.3% w/v monosaccharide. Agar (1.5 g) was added and the liquid was autoclaved, then cooled to approximately 50-60°C, before being poured into petri dishes, and allowed to cool to r.t. Frozen stocks of bacterial strains (*E. coli* ATCC 25922 or *Salmonella enterica* Typhimurium ATCC 14028) were thawed on ice, streaked on the agar plates, and the plates were incubated overnight at 37°C.

### Biomass hydrolysates for liquid cultures of *Paraburkholderia sacchari*

*Paraburkholderia sacchari* IPT 101 (DSM 17165) was purchased from the German Collection of Microorganisms and Cell Cultures (DSMZ, Germany). The strain was resuscitated in Reasoner’s 2A (R2A)^4^ medium (0.50 g/L yeast extract, 0.50 g/L proteose peptone (Difco no. 3), 0.50 g/L casamino acids, 0.50 g/L glucose, 0.50 g/L soluble starch, 0.30 g/L Na-pyruvate, 0.30 g/L K_2_HPO_4_, 0.50 g/L MgSO_4_.7H_2_O, final pH 7.2) as recommended by DSMZ and stored at −70°C.

The growth medium composition was adapted from a previous study.^5^ In brief, the nitrogen-limited medium (before carbon source addition) was composed of 1 g/L (NH_4_)_2_SO_4_, 6.78 g/L Na_2_HPO_4_.7H_2_O, 1.5 g/L KH_2_PO_4_, 1 g/L yeast extract, 0.2 g/L MgSO_4_, and 1 mL/L of trace elements solution (see agar plate section above).^6^ The pH was adjusted to 6.8. The MgSO_4_ stock solution (X100) was autoclaved and the trace elements stock solution (X1,000) was sterilized by filtration (0.22 μm). The remaining components were sterilized by autoclaving.

Carbon source stocks were prepared either from commercial glucose or diluted biomass hydrolysate. The glucose solution (80 g/L in growth medium) was prepared from pure glucose, purchased from Sigma Aldrich and sterilized by filtration (0.22 μm). The hydrolysate was suspended in culture medium to obtain 46.2 g/L glucose and 22.6 g/L xylose, and homogenized by vortexing. The supernatant was recovered after centrifugation (5 min, 4,700 × *g*), and sterilized by filtration (0.22 μm).

The bacterium from frozen stocks was first streaked on agar plates prepared from R2A medium before incubation at 30 C for 18 hours. One colony was used to inoculate the seeding medium (100 mL) in 250 mL shake flasks. The seeding medium had the same composition as the culture medium, except for a lower sugar concentration (10 g/L glucose and 5 g/L xylose). Following inoculation, the seeding medium was incubated at 30°C and 170 rpm for 18 hours to approximately 0.5 g/L cell dry mass (CDM). An aliquot of the seeding culture (1 mL) was transferred to the growth medium (15 mL) with the carbon source stocks (5 mL) or growth medium (5 mL, blank) and incubated at 30°C and 170 rpm for 24 hours.

Bacterial growth medium aliquots were taken and centrifuged for 5 min at 4,700 × *g*. The supernatant was filtered on 0.22 μm filter, and the filtrate was injected in the sugar analyzer.

### Cell dry mass determination

The cell dry mass was determined gravimetrically. In brief, 1.5 mL of culture broth was centrifuged (15,900 × *g* rpm for 5 min). The pellet was washed twice with distilled water (2 × 1 mL) and frozen-dried until constant weight (ca. 24 hours).

## Acknowledgements

We thank Iogen Corporation (Ottawa, Ont., Canada) for providing characterized WS and SB substrates, and POET (Sioux Falls, SD, USA) for providing characterized CS. We acknowledge funding from GreenCentre Canada, FRQNT (PR-254169), the Centre in Green Chemistry and Catalysis (FRQNT-2020-RS4-265155-CCVC), NSERC DG (RGPIN-2017-04107 and RGPIN-2017-Memorial Fellowship (SMFSU 507347-17).

## Footnotes

1 A. Sluiter, B. Hames, D. Hyman, C. Payne, R. Ruiz, C. Scarlata, J. Sluiter, D. Templeton, J. Wolfe *NREL Technical report* NREL/TP-510-42621 “Biomass and Total Dissolved Solids in Liquid Process Samples”, **2008**.

2 T. K. Ghose, *Pure Appl. Chem.* **1987**, *59*, 257-268.

3 T. B. Causey, S. Zhou, K. T. Shanmugam, L. O. Ingram, *Proc. Natl. Acad. Sci. U. S. A.* **2003**, *100*, 825-32.

4 https://www.dsmz.de/microorganisms/medium/pdf/DSMZ_Medium830.pdfs

5 M. T. Cesário, R. S. Raposo, M. C. M. D.de Almeida, F. van Keulen, B. S. Ferreira, M. M. R. da Fonseca, *New Biotechnol.* **2014**, *31*, 104-113.

6 B.S. Kim, S.C. Lee, S.Y. Lee, H.N. Chang, Y.K. Chang, S.I. Woo, *Biotechnol. Bioeng.* **1994**, *43*, 892-898.

‡ Enzyme mass was taken into account only when it was loaded as a solid (T. longibrachiatum). The CTec2 cellulases mix was purchased as a solution, and the liquid volume measured was used in the calculation of the *η* parameter.

§ calculated based on the average molecular weight of glucose and xylose theoretically released after quantitative hydrolysis of cellulose and xylan in CS, *i.e*. 167.6 g.mol^−1^.

