## Supporting Information for "Rapid mechanoenzymatic saccharification of lignocellulosic biomass without bulk water or chemical pre-treatment"

### Supplementary Information

Fabien Hammerer, Shaghayegh Ostadjoo, Karolin Dietrich, Marie-Josée Dumont, Luis F. Del Rio, Tomislav Friščić* and Karine Auclair*

### Table of Contents

1) Supplementary figures ........................................................................................................ S2
2) Supplementary tables .......................................................................................................... S8

### 1) Supplementary figures

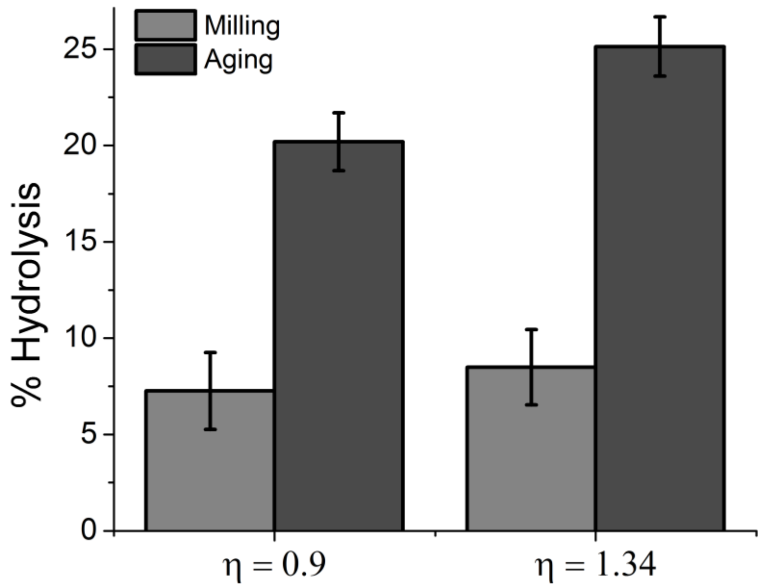


**Figure S1:** Effect of water loading on milling and aging reactions of WS (*η* in mg/mL. Pre-milled substrate (400 mg, 15 min at 30 Hz in a 15 mL s.s. jar with two 7 mm s.s. balls) was combined with *T. longibrachiatum* enzyme preparation (45.4 mg, 86 mg protein/g cellulose) and water (400 or 600 µL). The mixtures were milled at 30 Hz for 30 min and aged at 55°C for 3 days. Yield is DNS-based; error bars are standard deviation with n = 3.

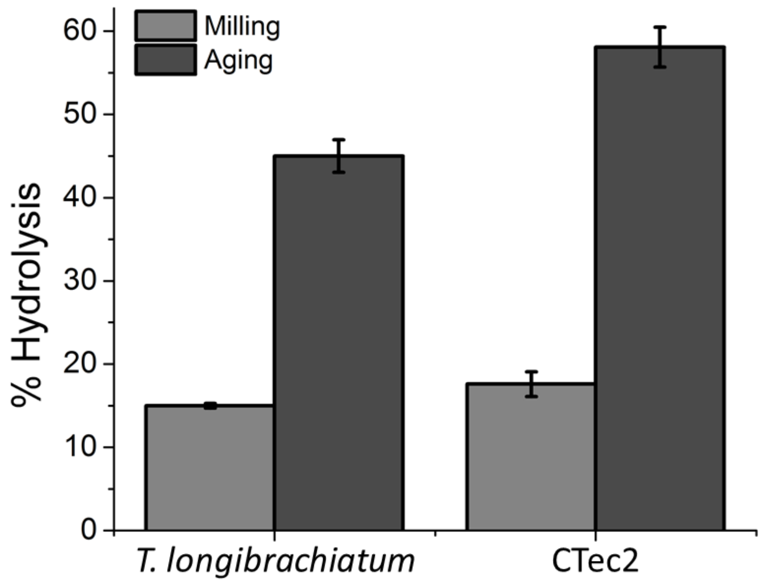


**Figure S2:** Mechano-enzymatic digestion of MCC (200 mg) at *η* = 1.0 µg/mL with *T. longibrachiatum* enzyme preparation (100 mg, 68 mg protein/g cellulose) or *CTec2* (57 µL, 45 mg protein/g cellulose). The mixtures were milled at 30 Hz for 30 min in a 15 mL s.s. jar with two 7 mm s.s. balls and aged at 55°C for 3 days. Yield is DNS-based; error bars are standard deviation with n = 3.

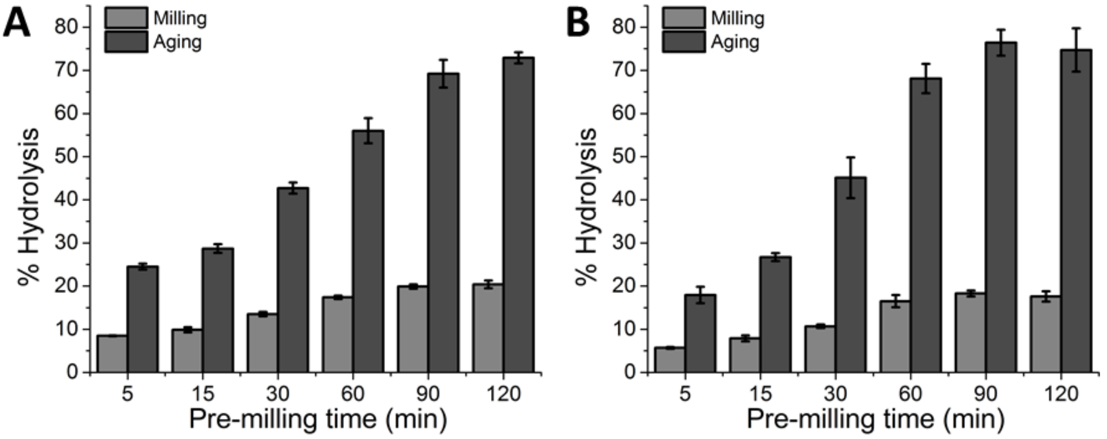


**Figure S3:** Influence of pre-milling time (3 g in 30 mL s.s. jar, 30 Hz) on the outcome of milling and aging reactions of WS (A) and SB (B). Substrate was combined with *CTec2* enzymes (45 mg/g cellulose) as a solution in water (600 µL, *η* = 1.5 µL/mg), milled 30 min at 30 Hz, aged 3 d at 55°C). Yield is DNS-based; error bars are standard deviation with n = 3.

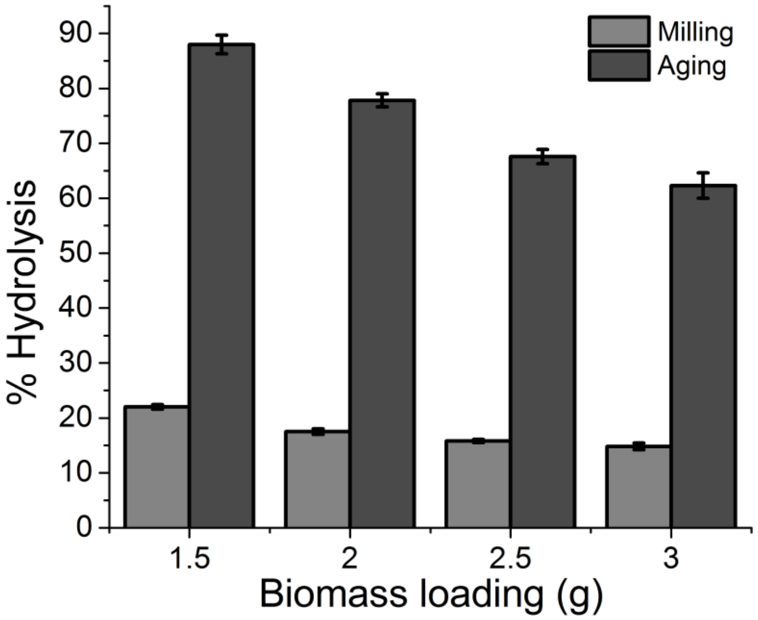


**Figure S4:** Influence of biomass loading on pre-milling efficacy of CS (60 min, 30 Hz) followed by milling and aging reactions with *CTec2* enzymes (45 mg/g cellulose) as a solution in water (600 µL, *η* = 1.5 µL/mg), milled 30 min at 30 Hz, aged 3 d at 55°C). Yield is DNS-based; error bars are standard deviation with n = 3.

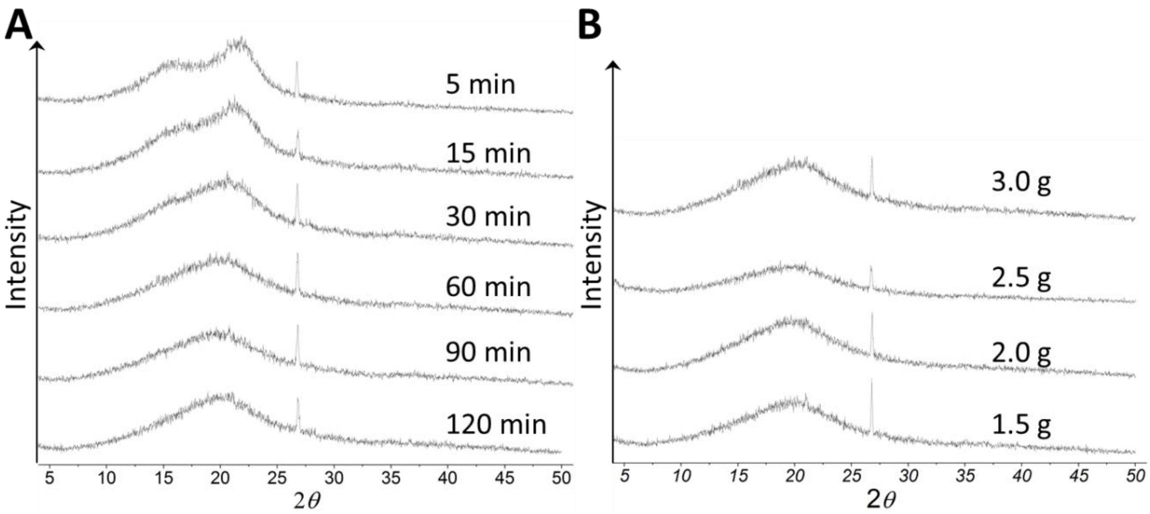


**Figure S5:** Comparison of PXRD patterns for CS after pre-milling. A) CS (3 g) pre-milled for various durations. B) CS pre-milled for 60 min at various loadings. Milling took place at 30 Hz in a 30 mL s.s. with one 15 mm s.s. ball. The crystalline cellulose peak at 22° disappears as ball-milling becomes longer and harsher. Peak at 27° is probably steel contamination.

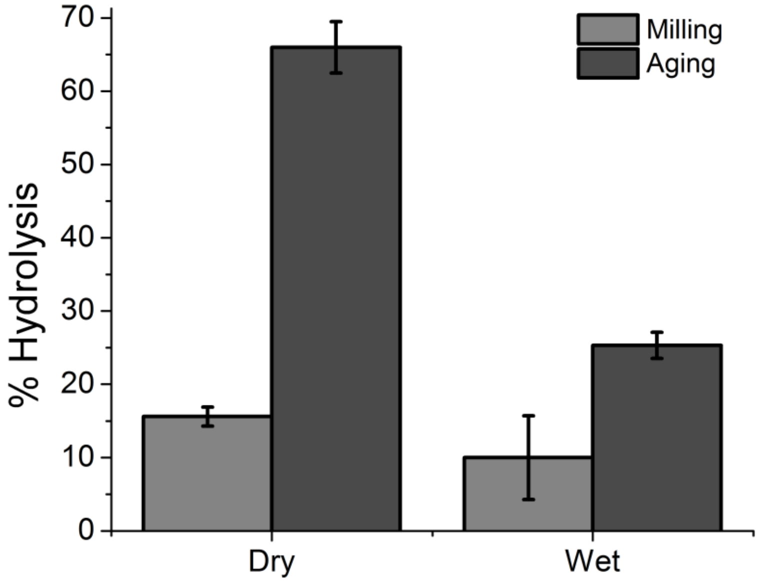


**Figure S6:** Influence of substrate humidity during pre-milling on the outcome of milling and aging reactions of CS. Substrate (400 mg dry i.e. 1.1% w/w water or 570 mg wet i.e. 30% w/w) was pre-milled for 30 min at 30 Hz in a 15 mL s.s. jar with two 7 mm s.s. balls. The resulting powder was then combined with *CTec2* enzymes (45 mg/g cellulose) as a solution in water (600 µL if dry or 430 µL if wet, $\eta$ = 1.5 µL/mg), milled 30 min at 30 Hz, and aged 3 d at 55°C). Yield is DNS-based; error bars are standard deviation with n = 3.

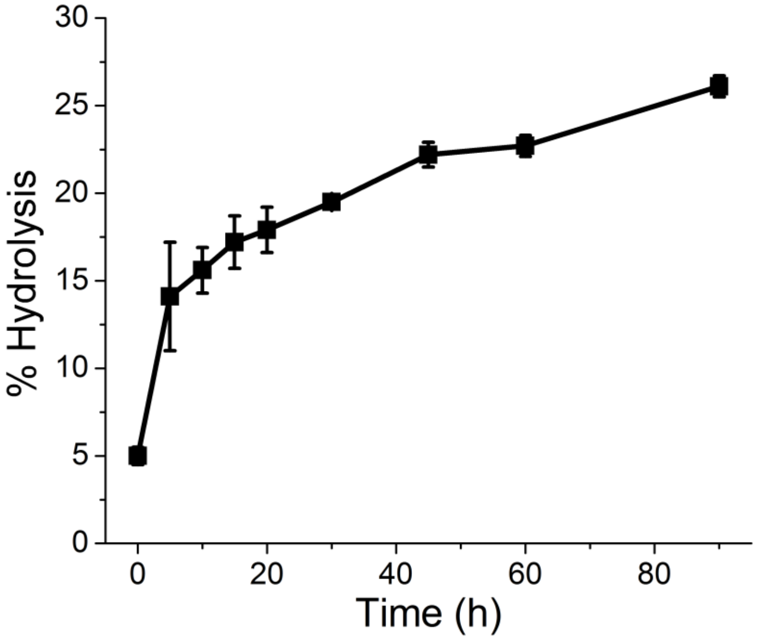


**Figure S7:** Kinetics of the hydrolysis reaction during the milling phase. CS (400 mg) was pre-milled (1.5 g, 60 min at 30 Hz in a 30 mL s.s. jar with one 15 mm s.s. ball) was combined with *CTec2* enzymes (45 mg/g cellulose) as a solution in water (600 µL, $\eta$ = 1.5 µL/mg) and milled at 30 Hz for 90 min. Yield is DNS-based; error bars are standard deviation with n = 3.

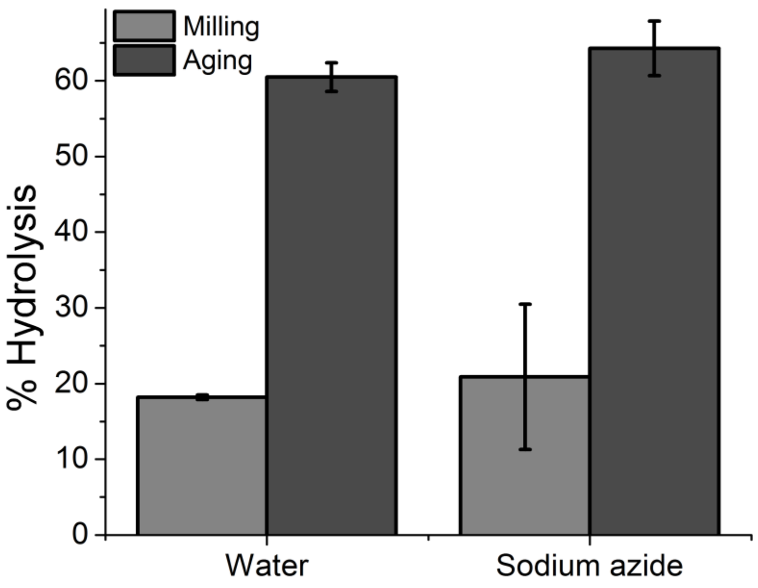


**Figure S8:** Influence of sodium azide on the outcome of milling and aging reactions. CS (400 mg) pre-milled (1.5 g, 60 min at 30 Hz in a 30 mL s.s. jar with one 15 mm s.s. ball) was combined with *CTec2* enzymes (45 mg/g cellulose) as a solution in water or $NaN_3$ 0.04% w/v (600 µL, $\eta$ = 1.5 µL/mg), milled 30 min at 30 Hz, and aged 3 d at 55°C). Yield is DNS-based; error bars are standard deviation with n = 3.

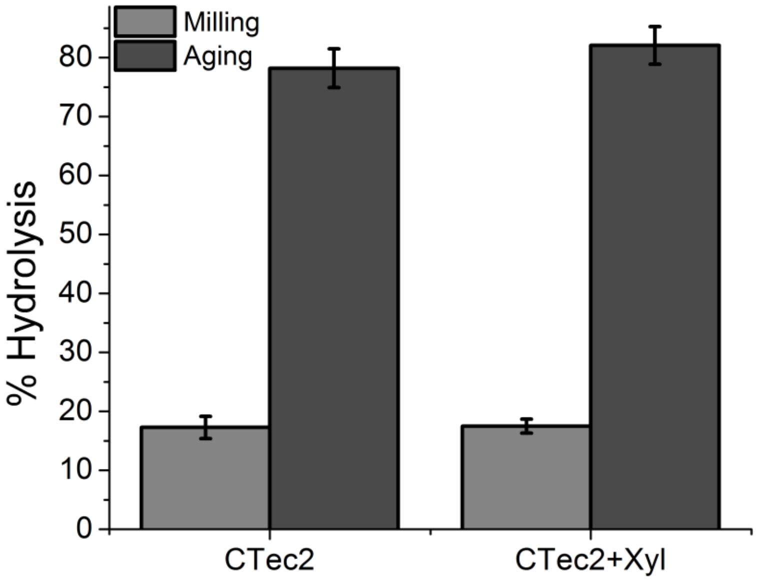


**Figure S9:** Influence of xylanase addition on the outcome of milling and aging reactions. CS (400 mg) pre-milled (1.5 g, 60 min at 30 Hz in a 30 mL s.s. jar with one 15 mm s.s. ball) was combined with *CTec2* enzymes (45 mg/g cellulose) as a solution in $NaN_3$ 0.04% w/v (600 µL, $\eta$ = 1.5 µL/mg) and 50 mg xylanase preparation from *Thermomyces lanuginosus* (1.5 mg/g cellulose), milled for 30 min at 30 Hz, and aged 3 d at 55°C. Yield is DNS-based; error bars are standard deviation with n = 3.

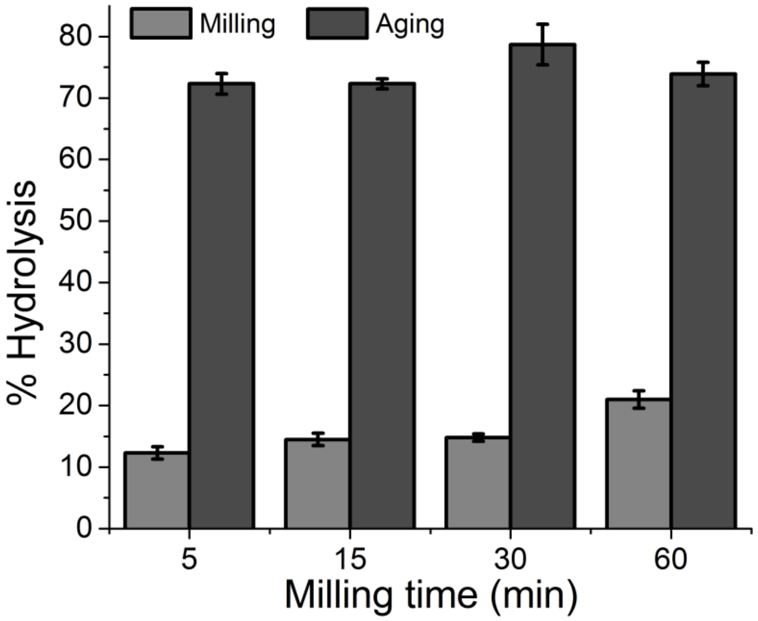


**Figure S10:** Influence of milling time on the outcome of milling and aging reactions. CS (400 mg) pre-milled (1.5 g, 60 min at 30 Hz in a 30 mL s.s. jar with one 15 mm s.s. ball) was combined with *CTec2* enzymes (45 mg/g cellulose) as a solution in $NaN_3$ 0.04% w/v (600 µL, $\eta$ = 1.5 µL/mg), milled at 30 Hz, and aged 3 d at 55°C). Yield is DNS-based; error bars are standard deviation with n = 3.

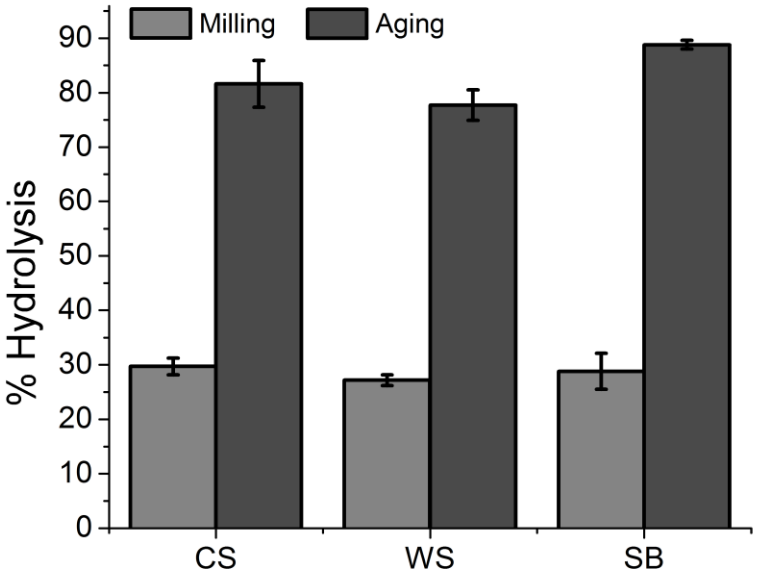


**Figure S11:** Scaling up of milling and aging reactions. Substrate (1.5 g) was pre-milled (60 min at 30 Hz in a 30 mL s.s. jar with one 15 mm s.s. ball) and combined with *CTec2* enzymes (45 mg/g cellulose) as a solution in $NaN_3$ 0.04% w/v (2.25 mL, $\eta$ = 1.5 µL/mg), milled for 5 min at 30 Hz, and aged 3 d at 55°C. Yield is DNS-based; error bars are standard deviation with n = 3.

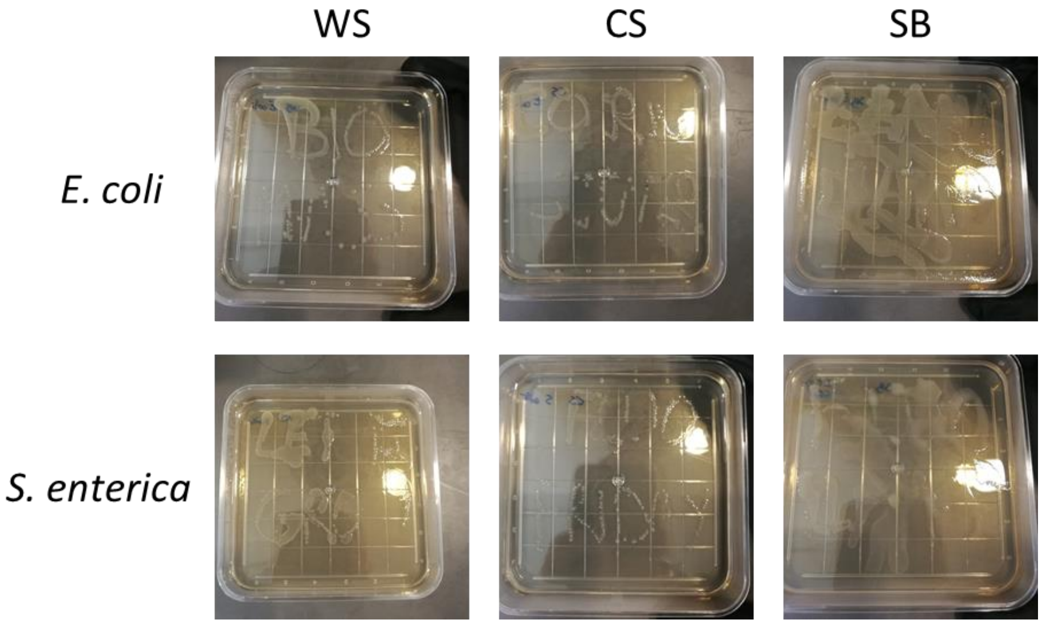


**Figure S12:** Bacterial (*E. coli* ATCC 25922 and *S. enterica* ATCC 14028) culture on media made from hydrolysates from CS, WS, and SB.

### 2) Supplementary tables

**Table S1.** Comparison of published processes for the digestion of cellulose in lignocellulosic biomass.

| Ref. | Substrate | Pre-treatment | Cellulases | Solid loading (g/L) | Enzyme loading[a] (FPU/g glucan) | Time (h) | Cellulose Conversion (%) | [Glucose][b] (g/L) | Productivity[c] (mg glucose/FPU) | Efficacy[d] (mg glucose/FPU/h) | Space-time yield[e] (g glucose/L/h) |
|---|---|---|---|---|---|---|---|---|---|---|---|
| | Corn stover[f] | Shaker milling (60 min) | CTec1 | 667 | 35[h] | 72 | 58 | 143 | 18.4 | 0.26 | 5.97 |
| | Corn stover[g] | Shaker milling (60 min) | CTec2 | 667 | 35[h] | 12 | 77 | 189 | 24.4 | 2.04 | 47.55 |
| | Sugarcane Bagasse[g] | Shaker milling (60 min) | CTec2 | 667 | 35[h] | 12 | 66 | 184 | 21.0 | 1.75 | 40.76 |
| | Wheat Straw[g] | Shaker milling (60 min) | CTec2 | 667 | 35[h] | 12 | 65 | 154 | 20.6 | 1.72 | 40.14 |
| 32 | Corn crop residues | Bioextrusion | ACCELERASE DUET | 400 | 58 | 48 | 11 | 22 | 2.1 | 0.04 | 1.02 |
| 32 | Corn crop residues | Bioextrusion | ACCELERASE DUET | 300 | 58 | 48 | 15 | 22 | 2.9 | 0.06 | 1.04 |
| 32 | Corn crop residues | Dilute base, bioextrusion | ACCELERASE DUET | 300 | 58 | 48 | 33 | 91 | 6.3 | 0.13 | 2.29 |
| 29 | Wheat straw | Hydrothermal (195°C, 19 min) | CTec2 | 300 | 14 | 72 | 60 | 94 | 47.6 | 0.66 | 2.78 |
| 56 | Corn stover | Steam explosion | Accelerase 1000 | 300 | 92 | 12 | 74 | 80 | 8.9 | 0.74 | 20.55 |
| 57 | Agave bagasse | Blade milling, hydrothermal (194°C, 30 min) | CTec2 | 250 | 15 | 72 | 98 | 196 | 72.6 | 1.01 | 3.78 |
| 58 | Rice straw | Dilute base, 24h | *T. reesei* + *A. niger* BG[l] | 250 | 18 | 48 | 56 | 155 | 34.6 | 0.72 | 3.24 |
| 59 | Corncob | Alkali liquor (12h, 70°C) | Cellulase A1 + Xylanase B1 | 200 | 40 | 120 | 41 | 57 | 11.4 | 0.09 | 0.76 |
| 60 | Rice straw | Dilute sulfuric acid (162°C, 10 min) | SacchariSEB-6 | 200 | 9 | 24 | 64 | 125 | 79.0 | 3.29 | 5.93 |
| 61 | Sugarcane Bagasse | Ambient glycerol organosolv (240°C, 30 min) | CTec2 + BG[i] + Endoxylanase + LPMO[j] AA9 | 200 | 3 | 72 | 83 | 105 | 307.4 | 4.27 | 2.56 |
| 62 | Brewers spent grains | Dilute HCl (121°C, 30 min) | CTec2 | 150 | 10[k] | 72 | 44 | 45 | 48.9 | 0.68 | 1.02 |
| 63 | Sugarcane Bagasse | Dilute alkaline (120°C, 60 min) | *C cubensis*, *P. pinophilum* | 120 | 34 | 120 | 51 | 40 | 16.7 | 0.14 | 0.57 |
| 64 | Wheat Straw | Oxygen delignification (10% NaOH, 150°C, 60 min) | Celluclast 1.5L + Novozyme 188 | 50 | 40 | 24 | 81 | 25 | 22.5 | 0.94 | 1.87 |

a) reported as Filter Paper Unit (FPU) per gram of cellulose. b) Final glucose concentration measured *in situ*. c) defined as the mass of glucose produced per enzymatic activity unit. d) defined as the mass of glucose produced per enzymatic activity per hour. e) defined as the mass of glucose produced per volume unit per hour. f) Result from aging reactions. g) Result from RAging reactions. h) extrapolated from Cannella et al. who used the same commercial enzyme cocktail and reported both protein titer and FPU activity. i) BG: β-glucosidase. j) LPMO: Lipid polysaccharide monooxygenase k) reported as FPU per gram of substrate.

**Table S2.** Comparison of published processes for the digestion of holocellulose (cellulose and hemicellulose) in lignocellulosic biomass.

| Ref. | Substrate | Pre-treatment | Cellulases | Solid loading (g/L) | Enzyme loading[a] (FPU/g glucan) | Time (h) | Holocellulose Conversion (%) | [RS][b] (g/L) | Productivity[c] (mg RS/FPU) | Efficacy[d] (mg RS/FPU/h) | Space-time yield[e] (g RS/L/h) |
|---|---|---|---|---|---|---|---|---|---|---|---|
| | Corn Stover[f] | Ball-milling (shaker, 60 min) | CTec2 | 667 | 16[h] | 24 | 69 | 306 | 47.9 | 2.00 | 21.30 |
| | Corn stover[g] | Ball-milling (shaker, 60 min) | CTec2 | 667 | 35[h] | 12 | 83 | 377 | 26.3 | 2.20 | 51.26 |
| | Sugarcane Bagasse[g] | Ball-milling (shaker, 60 min) | CTec2 | 667 | 35[h] | 12 | 76 | 353 | 24.1 | 2.01 | 46.93 |
| | Wheat Straw[g] | Ball-milling (shaker, 60 min) | CTec2 | 667 | 35[h] | 12 | 70 | 290 | 22.2 | 1.85 | 43.23 |
| 32 | Corn crop residues | Bioextrusion | ACCELERASE DUET | 400 | 58 | 48 | 20 | 62 | 3.8 | 0.08 | 1.85 |
| 32 | Corn crop residues | Bioextrusion | ACCELERASE DUET | 300 | 58 | 48 | 25 | 58 | 4.8 | 0.10 | 1.74 |
| 32 | Corn crop residues | Dilute base, bioextrusion | ACCELERASE DUET | 300 | 58 | 48 | 42 | 97 | 8.0 | 0.17 | 2.92 |
| 58 | Rice straw | Dilute base, 24h | *T. reesei* + *A. niger* BG[i] | 250 | 18 | 48 | 73 | 203 | 45.1 | 0.94 | 4.22 |
| 59 | Corncob | Alkali liquor (12h, 70°C) | Cellulase A1 + Xylanase B1 | 200 | 40 | 120 | 43 | 103 | 11.9 | 0.10 | 0.80 |
| 60 | Rice straw | Dilute sulfuric acid (162°C, 10 min) | SacchariSEB-6 | 200 | 9 | 24 | 68 | 133 | 83.9 | 3.50 | 6.30 |
| 61 | Sugarcane Bagasse | Ambient glycerol organosolv (240°C, 30 min) | CTec2 + BG[i] + Endoxylanase + LPMO[j] AA9 | 200 | 3 | 72 | 94 | 155 | 348.1 | 4.83 | 2.90 |
| 63 | Sugarcane Bagasse | Dilute alkaline (120°C, 60 min) | *C. cubensis*, *P. pinophilum* | 120 | 34 | 120 | 58 | 63 | 19.0 | 0.16 | 0.64 |
| 64 | Wheat Straw | Oxygen delignification (10% NaOH, 150°C, 60 min) | Celluclast 1.5L + Novozyme 188 | 50 | 40 | 48 | 84 | 37 | 23.3 | 0.49 | 0.97 |

a) reported as Filter Paper Unit (FPU) per gram of cellulose. b) Final reducing sugar (RS) concentration measured *in situ*. c) defined as the mass of reducing sugar produced per enzymatic activity unit. d) defined as the mass of reducing sugar produced per enzymatic activity per hour. e) defined as the mass of reducing sugar produced per volume unit per hour. f) Result from aging reactions. g) Result from RAging reactions. h) extrapolated from Cannella et al. who used the same commercial enzyme cocktail and reported both protein titer and FPU activity. i) BG: β-glucosidase. j) LPMO: Lipid polysaccharide monooxygenase
